# A report card methodology to showcase progress towards threatened species recovery

**DOI:** 10.1101/2022.09.06.506545

**Authors:** Michelle Ward, Tracy Rout, Romola Stewart, Hugh P. Possingham, Eve McDonald-Madden, Thomas G. Clark, Gareth S. Kindler, Leonie Valentine, Ellen Macmillan, James E.M. Watson

**Affiliations:** WWF-Australia, Level 4B, 340 Adelaide Street, Brisbane QLD 4000; Centre for Biodiversity and Conservation Science, The University of Queensland, St Lucia, Queensland, Australia

**Keywords:** Australia, biodiversity, conservation, policy, reporting, extinction

## Abstract

Among the conservation community, it is well known that Earth’s mass species extinction crisis is getting worse. Yet, an often neglected problem is the need for effectively communicating the species extinction crisis to diverse audiences in ways that catalyse immediate action. Here we generated a streamlined threatened species recovery report card methodology, which combined two input indicators including planning and funding, one output indicator capturing habitat protection, and one outcome indicator which highlights threatened species trajectories, to provide simple scores for all Australian threatened species. We show that just 41 (2.3%) of species achieved an A grade for the input indicator (i.e., recovery plans and federal funding), 240 (13.3%) achieved a C grade, and 1,521 (84.4%) achieved an F. Five hundred and twenty nine (29.3%) species achieved an A for the output indicator (i.e, habitat protection), 130 (7.2%) achieved a B, 158 (8.8%) achieved a C, 189 (10.5%) achieved a D, 212 (11.8%) achieved an E, and 584 (32.4%) achieved a F. While five (0.3%) species achieved an A for the output indicator (i.e., threat status improvement), every other species (99.7%) achieved an F. We provide a method to combine scores to test how individual jurisdictions are tracking and show that Australia is achieving an F for the input and outcome indicators, and a D for the output indicator. While the threatened species recovery report card highlighted a clear failure in many federal environmental legislation responsibilities, it provides a baseline from which different governments can track policy progress and outlines clear direction for immediate improvement including developing adequate recovery plans, funding the actions in the recovery plans, protecting habitat from further destruction, verifying recovery through monitoring and evaluation of species trajectories, and supporting transparency and collaboration on the execution on the plans through an improved data infrastructure. Without an immediate step change in how Australia communicates and faces its species crisis, we will leave a tragic legacy of extinction and fail our obligations to future generations of Australians, and the international community.

## Introduction

Earth is currently experiencing a sixth mass species extinction event driven by human activities (Johnson et al. 2017). It has been estimated that millions of species are now well on the pathway to extinction, affecting the functioning of all ecosystems and the essential services they provide to humanity (IPBES 2019). Biodiversity loss is now considered a serious threat to humanities’ long-term well-being (Díaz et al. 2019) and nations have responded by setting ambitious conservation targets to reverse loss through various mechanisms, including the United Nation’s Sustainable Development Goals (SDGs) and Convention on Biological Diversity (CBD) (United Nations Sustainable Development Goals 2015; Secretariat of the Convention on Biological Diversity 2021).

It is well established that abating species extinction and achieving species recovery in-situ is often complicated and multifaceted (Akçakaya et al. 2018; Stephenson et al. 2020). For success to occur, it often needs proactive action, at different spatial scales, by numerous actors, including scientists, practitioners, governments, and landholders who may have to embrace activities they have not done in the past or are not in their best financial interest (Watson et al. 2022). But beyond the practicalities of achieving a positive outcome, an often under-appreciated, but increasingly important part of securing threatened species is communicating the extent of the problem to mobilise action (Novacek 2008b). Crucially, this communication (on what is needed to secure species) needs to be done in ways that the wider community understands and relates to, thus galvanising the support needed to catalyse action.

While significant exemplary efforts have been made to document the size of the threatened species problem in global and nation reports (Convention on Biological Diversity 2011; IPBES 2019; Secretariat of the Convention on Biological Diversity 2020; WWF 2020), the limited public knowledge of the growing extent of the biodiversity crisis globally (beyond concerned citizens and experts) is a recognised problem (Novacek 2008a; Natural History Museum 2020). It is argued that the Anthropocene continues to be a period of “unprecedented looking away” by people when considering their impact on the environment (Haraway 2016). With millions of species declining at ever faster rates (Ceballos et al. 2015), it is evident that clearer communication on the extent of the global species extinction crisis is required. Communication of the biodiversity crisis must be engaging and localised for greater impact (Gobby et al. 2021), as well as provide opportunities for stronger stewardship when relevant (Rousell & Cutter-Mackenzie-Knowles 2020).

Here, we develop a broad methodology to generate a scalable threatened species recovery report card with the aim that it can be used to communicate the state of any threatened species to the wider community in an easy-to-understand format. It can be aggregated to report across any scale of jurisdiction (i.e., from a single protected area, local government area, electorate, state, to a nation or continent) to provide an easy reporting tool across time to assess progress. The report card goes beyond providing information on the actual state of individual species (such as those efforts provided by IUCN Red List of Threatened Species, (IUCN 2018)) as it utilises a input-output-outcome framework (Margoluis et al. 2013) to highlight information on conservation progress. The report card contains known information on levels of planning, funding, and protection, and recent trajectories for each species, which when combined, provides a simple communication tool the state of each species. It builds on efforts of the IUCN Green List of Species (Akçakaya et al. 2018) by being a communication tool for conservation progress rather than a planning and reporting apparatus for species’ ecological recovery.

We showcase this reporting method by utilising data on threatened species in Australia. As one of just 17 mega-biodiverse nations (Commonwealth of Australia, 2022), Australia has one of the worst records for recent species extinctions (Ritchie et al. 2013), and it is likely many more will go extinct without significant investment in proactive actions (Garnett et al. 2022; Kearney et al. 2022), making it a good case study. Australia is also a signatory to the Convention of Biological Diversity (CBD)and Sustainable Development Goals (SDGs; Secretariat of the Convention on Biological Diversity, 2020) and has clear obligations under national environmental legislation for protecting threatened species (e.g., Environment Protection and Biodiversity Conservation Act; EPBC Act, (Commonwealth of Australia 1999)). We showcase the use of the report card over different scales by disaggregating threatened species data by national, electorate, state and territory, and local government areas (LGA), of which jurisdictions play different, but important roles in securing Australian threatened species.

## Method

The threatened species report card utilises the input-output-outcome framework to help visualise the relationship between resources used (inputs), activities undertaken (outputs), and the results (outcomes). This logic model is used because of its generality and easy to understand theory of change (Margoluis et al. 2013). While results chains are more specific and show direct relationships between inputs, outputs, and outcomes (Margoluis et al. 2013), the type of data needed for result chains is not available for most threatened species and while critical for recovery planning, is not needed as a communication tool, thus a simpler logic model was used.

The report card comprises of four indicators (two inputs, one outcome, and one output), covering the major building blocks needed to recover threatened species. The four indicators include known information on planning, funding, levels of protection, and recent trajectories for each species. These indicators are assessed against the desired state which is to ensure that all species have recovery plans (input), all species have dedicated funding to secure their management needs (input), habitat for all species has adequate protection (outcome), and species are improving in extinction risk trajectories (output). We chose these indicators not only because they are critical to recovering threatened species, but because they are measurable, data is often updated by the Australian government to ensure repeatability in future years and can be combined to provide a simple communication tool for species, and if needed, for any jurisdiction of interest (**Table 1**).

**Table 1.** Summary of the four indicators within the threatened species recovery report card for species and jurisdictional reporting

| **Indicators** | **Indicator for species reporting** | **Indicator for jurisdictional reporting** | **Method for species reporting** | **Method for jurisdictional reporting** | **Benchmark** | **Action** |
| --- | --- | --- | --- | --- | --- | --- |
| **Input: Recovery plans** | Species has a recovery plan. | Proportion of species with recovery plan. | Identify if species has a recovery plan made or updated in the last 10 years. | Number of species with recovery plan made or updated in the last 10 years divided by total number of species. | 1 = Species (or all species in the jurisdiction) has a recovery plan made/updated within the last 10 years. | Ensure the timely development and review of recovery plans. |
| **Input: Funding recovery** | Species has had dedicated federal government funding in the last 5 years. | Proportion of species with dedicated federal government funding in the last 5 years. | Identify species with dedicated funding. | Number of species with funding divided by total number of species. | 1 = Species (or all species in the jurisdiction) has received some dedicated funding in the last 5 years. | Fund actions in species' recovery plans. |
| **Output: Species-specific habitat protection** | Species habitat has enough protection to meet species-specific protection target. | Proportion of the species-specific protection target that has been achieved in threatened species habitat. | Spatially intersect all threatened species habitats with protected area estate. | The average of the species-specific protection target that has been achieved in threatened species habitat within the jurisdiction. | 1 = Species (or all species in the jurisdiction) has more than or equal to their species-specific habitat protection target. | Commit and implement adequate habitat protection for each species habitat. |
| **Outcome: Species extinction risk trajectories** | Species has improved in threat status over the last 10 years. | Proportion of species that have improved in threat status over the last 10 years. | Identify if species threat status has improved. | Number of species whose threat status has improved divided by the total number of species | 1 = Species (or all species in the jurisdiction) has improved in threat status in the last 10 years. | Evidence that species are recovering. |

### Input indicator: Recovery plans

Once a species has been listed as threatened with extinction, recovery plans are critical as they provide a roadmap to stopping the decline, and supporting the recovery of species (Department of the Interior 1973; Commonwealth of Australia 1999). Well-funded recovery plans ensure transparency and accountability through clear documentation of species threats, actions, and expenditure needed, along with monitoring and evaluation of conservation outcomes (Bottrill et al. 2011; McDonald et al. 2015; Wintle et al. 2019). A recent global analysis of threatened species found that more than half of species required species-specific recovery actions to prevent extinction, in addition to area-based conservation and threat management (Bolam et al. 2022). Without recovery plans for each species, necessary species-specific recovery actions may remain unidentified and recovery efforts ineffective.

### Input indicator: Funding recovery

Earth’s ongoing species loss and growing threatened species list is a direct result of inadequate funding for environmental protection and targeted species recovery. Research shows that the more a country spends on conservation, the fewer species extinctions (Waldron et al. 2017). In the United States, strong investment in threatened species recovery has resulted in 39 species being de-listed due to recovery efforts (Wintle et al. 2019). The critical need for adequate financial resources is exemplified in international commitments, such as the CBD where Goal D demands that the gap between available financial and other means of implementation, and those necessary to achieve the 2050 Vision of ‘living in harmony with nature’, is closed (Secretariat of the Convention on Biological Diversity 2021). To recover threatened species, there must be adequate investment in appropriate conservation actions.

### Output indicator: Protection of threatened species habitats

Protected areas are the cornerstone in biodiversity conservation (Watson et al. 2014a; Dudley & Stolton 2016; Maxwell et al. 2020). With threatened species habitat continually being destroyed and successive climate-driven conservation disasters, such as droughts, bushfires, and floods also taking a devastating toll (Ward et al. 2019, 2020; Legge et al. 2021), protected areas will continue to play a key role in protecting habitat and recovering threatened species (Watson et al. 2014b). When protected areas are connected, representative, and well managed, they offer a simple solution that safeguards habitat, builds resilience to climate change, and facilitates fundamental ecological processes such as migration and dispersal (Thomas & Gillingham 2015; Barr et al. 2016; Tucker et al. 2018).

### Outcome indicator: Species extinction risk trajectories

Under the IUCN Red List there are seven categories to classify extinction risk: least concern, near threatened, vulnerable, endangered, critically endangered, extinct in the wild, and extinct (IUCN 2015). These categories are generally consistent across countries with some variations. For example, under Australian federal listings, there is no least concern or near threatened category. This classification changes if a species becomes more or less threatened over time (Monroe et al. 2019). There is now extensive research that shows when conservation actions are well-planned, funded, and implemented, they can stop species extinction, slow the extinction rate, and halt and reverse species extinction risk trajectories (Mace et al. 2018; Monroe et al. 2019; Bolam et al. 2021). Thus, species extinction risk trajectories are important to capture and highlight successful recovery efforts.

All four indicators can be used to produce threatened species grades (**Fig. 1**). These grades can be utilised to communicate the state of any threatened species at any point in time. The indicators can also be aggregated to report across any scale of jurisdiction. These jurisdictions could include site scale, protected areas, local government areas, electorates, states, or even used for national reporting. Scores were converted to grades where an F grade is less than 0.16, an E grade is from 0.17-0.32, a D grade is 0.33 – 0.49, a C grade is 0.5 – 0.66, a B grade is 0.67 – 0.82, and an A grade is greater than 0.83.

**Figure 1.**
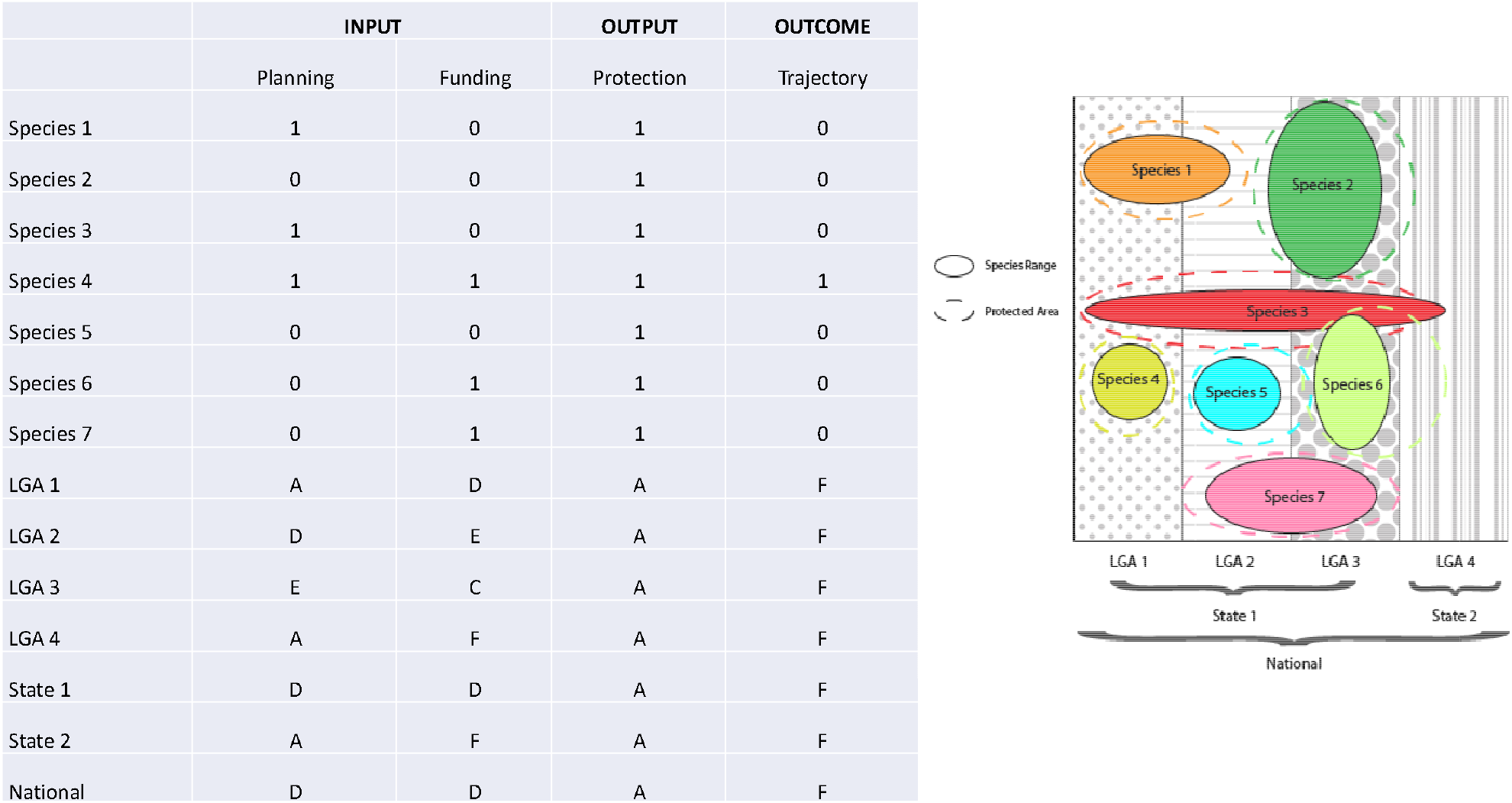
Method figure outlining the four indicators and how each were scored across multiple jurisdictions.

### Australia as a case study

As of 7^th^ March, under the EPBC Act, 1,948 species threatened with extinction were placed into one of six threat categories: Conservation dependent, Vulnerable, Endangered, Critically Endangered, Extinct in the wild, or Extinct. Except for Extinct species (n=105), most are mapped within the Australian Government’s publicly available Species of National Environmental Significance (SNES) spatial dataset (Commonwealth Government 2022) last updated March 2022. The SNES maps are modelled distributions of each threatened species that include known and likely areas of occurrence. The maps have two categories: areas where the species or species habitat is ‘likely to occur’, and areas where the species or species habitat ‘may occur’. The SNES maps are generalised to a 1km_2_ grid resolution, or a 10km_2_ resolution for sensitive species, therefore may overestimate where species occur (Commonwealth Government 2022). To refine this, we used only areas where the species or species habitat is ‘likely to occur’, as ‘may occur’ areas are too broad for this analysis. As we focus on threatened terrestrial and freshwater species, we removed 27 threatened marine species from the SNES spatial dataset; however, retained some semi-aquatic marine species, such as sea turtles, pinnipeds, and sea birds. To determine the occurrence of the remaining 1,816 threatened species in each state and territory, and local government area, we overlaid the SNES spatial dataset (Commonwealth Government 2022) with the State and Territory boundaries and local government areas to create species spatial matrices.

For species that have been delisted (n=5) in the last 10 years or were missing from the SNES spatial dataset (n=7), we sourced occurrence records using Atlas of Living Australia. These occurrence records were overlaid with the 2021 Federal Electoral Division maps (Australian Electoral Commission 2022). Capricorn Rabbit-rat (*Conilurus capricornensis*), Blue-grey Mouse (*Pseudomys glaucus*), Thalassarche bulleri platei, Diomedea antipodensis gibsoni, Leucocarbo atriceps nivalis, Leucocarbo atriceps purpurascens, Lomatia tasmanica, and South-eastern Striped Bandicoot (*Perameles notina*) had no occurrence records in the Atlas of Living Australia and were not included in the analysis.

### Recovery plans

When a species is listed under the EPBC Act, a Conservation Advice is developed which commonly details the species taxonomy, threat category listing advice, threats, and actions. Some species also have a recovery plan, which sets out the research and management needed to protect and recover species, as well as how to manage and reduce threatening processes. Recovery plans sometimes also outline the costs of expected actions and a plan for implementation including co-ordination of government agencies and stakeholders. While both are important conservation tools, the Minister for Environment is legally bound to act consistently with a recovery plan, while conservation advice documents are not legally enforceable.

For our assessment, we use recovery plans as the legally binding instrument as the best indicator for tracking federal government planning and action for recovery. This does not mean previously developed recovery plans are perfect. Rather, they have been criticised for taking too long to implement, vague wording, and not including key information such as costs or actions to mitigate threats such as habitat loss (Bottrill et al. 2011). Effective recovery plans must be concise and time-bound, include return on investment principles, and must include a plan for monitoring and evaluation of conservation outcomes (Bottrill et al. 2011). Most importantly, the actions outlined in recovery plans must be funded and implemented, preferably through long-term and flexible funding arrangements (Wintle et al. 2019).

Legally, recovery plans remain in force until their legislated sunsetting date, unless the species is de-listed. The legislated sunsetting date is the 1 April or the 1 October, 10 years after the recovery plan was made or adopted, although this date can be deferred by the minister (Commonwealth of Australia 2022a). We used publicly available data downloaded from the Australian Governments’ Species Profile and Threats (SPRAT) database (Commonwealth of Australia 2022a) to assess whether each threatened species has a current recovery plan, defined as a plan adopted after the 2nd October 2011 through to 7^th^ March 2022, to align with mapping dataset. For the nation, and for each electorate, state and territory, and LGA we summed the number of species with a current recovery plan, divided by the total number of species in the nation, electorate, state and territory, and LGA.

### Funding recovery

Currently, there is no centralised system for tracking the funding provided to each federally listed threatened species, nor is there any systematic way to track whether the actions identified and costed in recovery plans have been funded or implemented. Ideally, there would be costed recovery plans whereby one could report on adequate funding. In the absence of a properly managed funding data tracking system, to assess which threatened species have received funding for their recovery, we collated all publicly available information on Australian Government funded conservation projects in the last 5 years and included all projects regardless of amount or appropriateness to recovering species. This included projects funded through the Threatened Species Recovery Fund, the Environment Restoration Fund, the National Landcare Program Regional Land Partnerships, the Bushfire Recovery Fund Phase 1 and Phase 2, and the Koala Conservation Package (Supplementary Table 1). For each of these projects we identified threatened species that were named in the title or description of the project, and then consolidated these to compile a list of all threatened species that have received dedicated Australian Government funding in the last 5 years.

In addition to the sources mentioned above, we investigated yearly Portfolio budget statements but found these did not identify funding down to the project-level and so did not include information about targeted species. Similarly, currently available information on the Oceans Leadership Package and the Reef 2050 Long-Term Sustainability Plan did not identify targeted species. We calculated the indicator for the nation, and each electorate, state and territory, and LGA by summing the number of species that received dedicated funding in the last 5 years, divided by the total number of species in the nation and each electorate, state and territory, and LGA.

This indicator captures only conservation funding that targets threatened species and mentions those species by name. The Australian federal government and state governments also fund threat abatement actions, such as fox (*Vulpes vulpes*) or cat (*Felis Catus*) control, which likely benefit a suite of threatened species. However, if recovery plans for each threatened species were costed, funded, implemented, and monitored, then there would by necessity be dedicated reported funding for every threatened species. Therefore, while this indicator will underestimate the number of threatened species benefiting from conservation action funded by the Australian government, it is an accurate measure of how well the government is systematically funding, tracking, and reporting progress in recovering species.

### Habitat protection

To ensure persistence and recovery, species conservation plans often state a requirement to have a measurable target threshold to be conserved by protecting remnant habitat and/or restoring degraded sites (Andren 1994; Maron et al. 2012; Simmonds et al. 2019). This target-based approach is often based on the mathematical relationship between habitat area and the number of species an area can support, generally referred to as the “species– area curve” (e.g., MacArthur & Wilson 1967). In recognising that setting meaningful targets should be species-specific, we used methods developed by Watson et al. 2010 (based on (Rodrigues et al. 2004) where we set a target for protection of either 1,000 km_2_ or 100% of the range of the species range, whichever value was smaller for species with a geographic range of <10,000km2. For all species with a geographic range of >10,000 km2, the target was at 10% of the range. We note that many species distributions have substantially contracted since pre-European colonisation, so these targets are likely an underestimate of how much is actually needed to delist species from the EPBC Act threatened species list.

Following the methodology by Sutcliffe et al. 2015, we calculated the proportion of the protection target that has been achieved by dividing the percentage protected by the species-specific protection target. Then for the nation, electorates, states and territories, and LGAs, we calculated the mean proportion of the protection target that has been achieved across the intersecting species.

While we recognise that the degree of connectivity, size of habitat patches, and effective management is of crucial importance to the viability of some species recovery and persistence, due to the lack of species-specific data, these elements are not captured in this analysis.

### Species threat status trajectories

To calculate species threat status trajectories, we analysed publicly available data from the Species Profiles and Threats database (Commonwealth of Australia 2022a) on threatened species delistings or transfers between threat categories (e.g., Vulnerable, Endangered, or Critically endangered) since commencement of the EPBC Act. We categorised each downlisting transfer and verified a genuine recovery by referring to Conservation Advices.

To find species whose status has recently improved, we looked at species that have been downlisted in the last 10 years (on or after 1 January 2012). We calculated the indicator for the nation, and each electorate, state and territory, and LGA by summing the number of species whose status has recently improved divided by the total number of species in the nation, and each electorate, state and territory, and LGA. This data calculates the proportion of species that have improved over the last 10 years in the nation and per electorate, state and territory, and LGA.

### The final grades of the threatened species recovery report card

The indicators were averaged across input indicators (i.e., recovery plans and funding), output indicator (i.e., protection), and the outcome indicator (i.e., species threat trajectories) to provide overall species scores. We also calculated input, output, outcome, grades for national, electorate, state and territory, and local government area reporting. As delisted species do not have recovery plans, funding, or maps to quantify protection, these species were removed in the average input and output indicator scores.

## Results

Ideally, every threatened species would have a recovery plan, dedicated funding, adequate species-specific protection, which would then improve threat status. Using our new method for calculating a threatened species report card, our analysis showed that only 173 species (9.6%) have up-to-date recovery plans (54 = Vulnerable, 69 = Endangered, 50 = Critically endangered) and 151 (8.3%) with dedicated funding (36 = Vulnerable, 75 = Endangered, 38 = Critically endangered; **Fig 2**). Of the 151 species that were identified as receiving funding, more than half of these (88) were mentioned in a single project (Supplementary Table 1). Eleven species were named in more than 10 conservation projects, and one species (koala) was named in 38 projects (Supplementary information 1). While 1,702 have some protection, only 385 have adequate protection according to their species-specific target. In the last 10 years, there have only been five genuine improvements of species threat status (excluding humpback whale (*Megaptera novaeangliae*) which was not analysed here as it’s a marine species).

**Figure 2.**
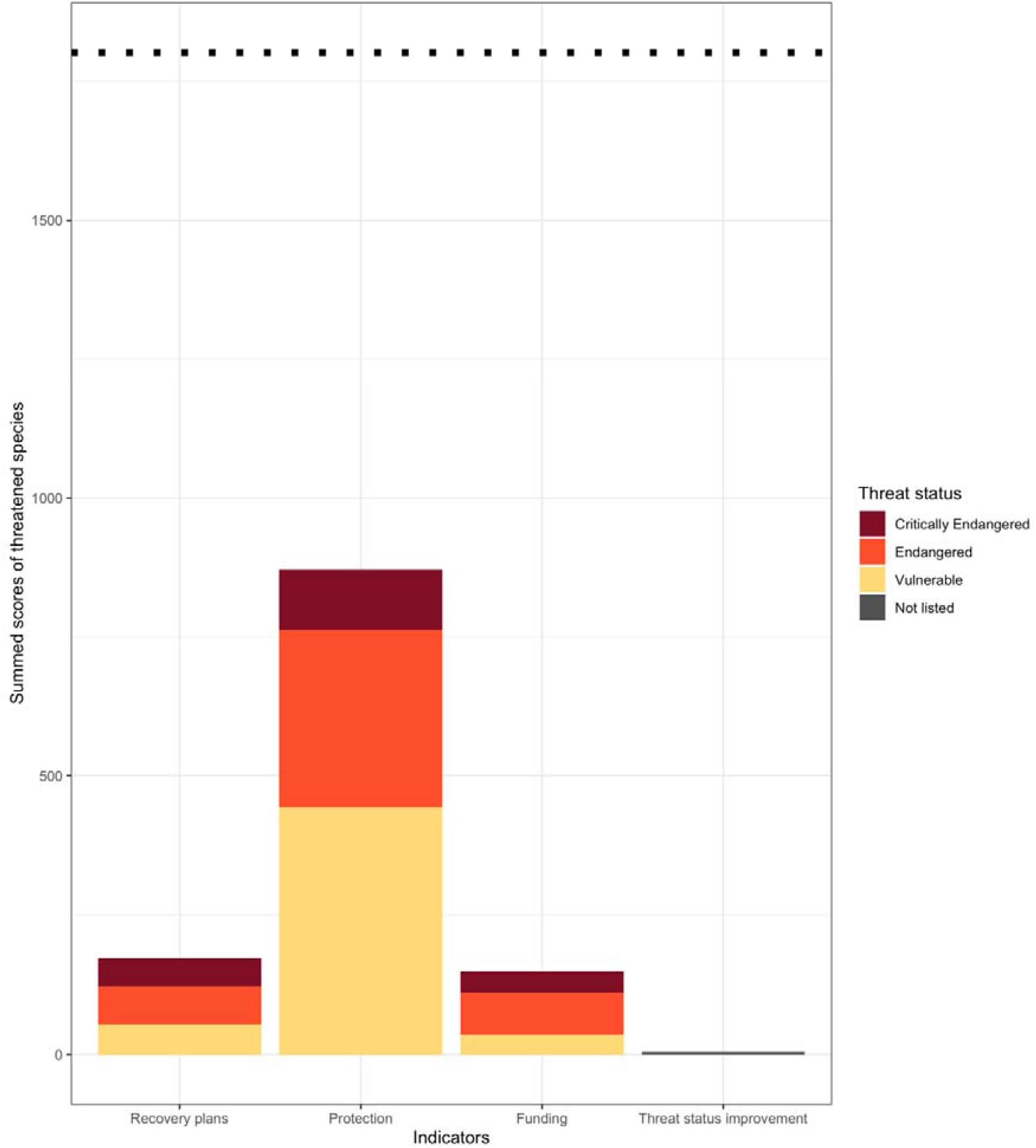
Summed scores of all threatened species in the analysis. The funding, recovery plans, and threat status improvement indicators are either 0 (no) or 1 (yes) for each species. Protection is a proportion of the species-specific target. The dotted line represents the benchmark target, highlighting the need for every species to have a recovery plan, funding, adequate protection, and threat status improvement. Critically Endangered species are highlighted maroon, Endangered species are highlighted orange, Vulnerable species are highlighted yellow, and Not listed species are highlighted grey.

When we averaged the two input indicators across every threatened species, we found that 41 species (2.3%) received an A grade, including spotted-tail quoll (*Dasyurus maculatus maculatus* (SE mainland population)), growling grass frog (*Litoria raniformis*), and regent honeyeater (*Anthochaera phrygia*), all of which have a recovery plan and dedicated funding (**Fig. 3A**). Two hundred and forty species (13.3%) received a C grade, and 1,521 species (84.3%) received a F grade. When we analysed the output indicator (i.e., protection), we found that 532 species (29.5%) received an A grade, 130 (7.2%) received a B grade, 156 (8.6%) received a C grade, 192 (10.6%) received a D grade, 218 (12.0%) received an E grade, and 574 (31.8%) received a F grade (**Fig. 3B**). Only five species (0.3%) received an A for the outcome indicator, all other species received an F (99.7%; **Fig. 3AC**). There were 122 (6.8%) species which did not have a recovery plan, funding, protection, and no threat status improvement. This included species such as Rottnest bee (*Hesperocolletes douglasi*), Tiwi masked owl (*Tyto novaehollandiae melvillensis*), and Mount Cooper striped skink (*Lerista vittata*).

**Figure 3.**
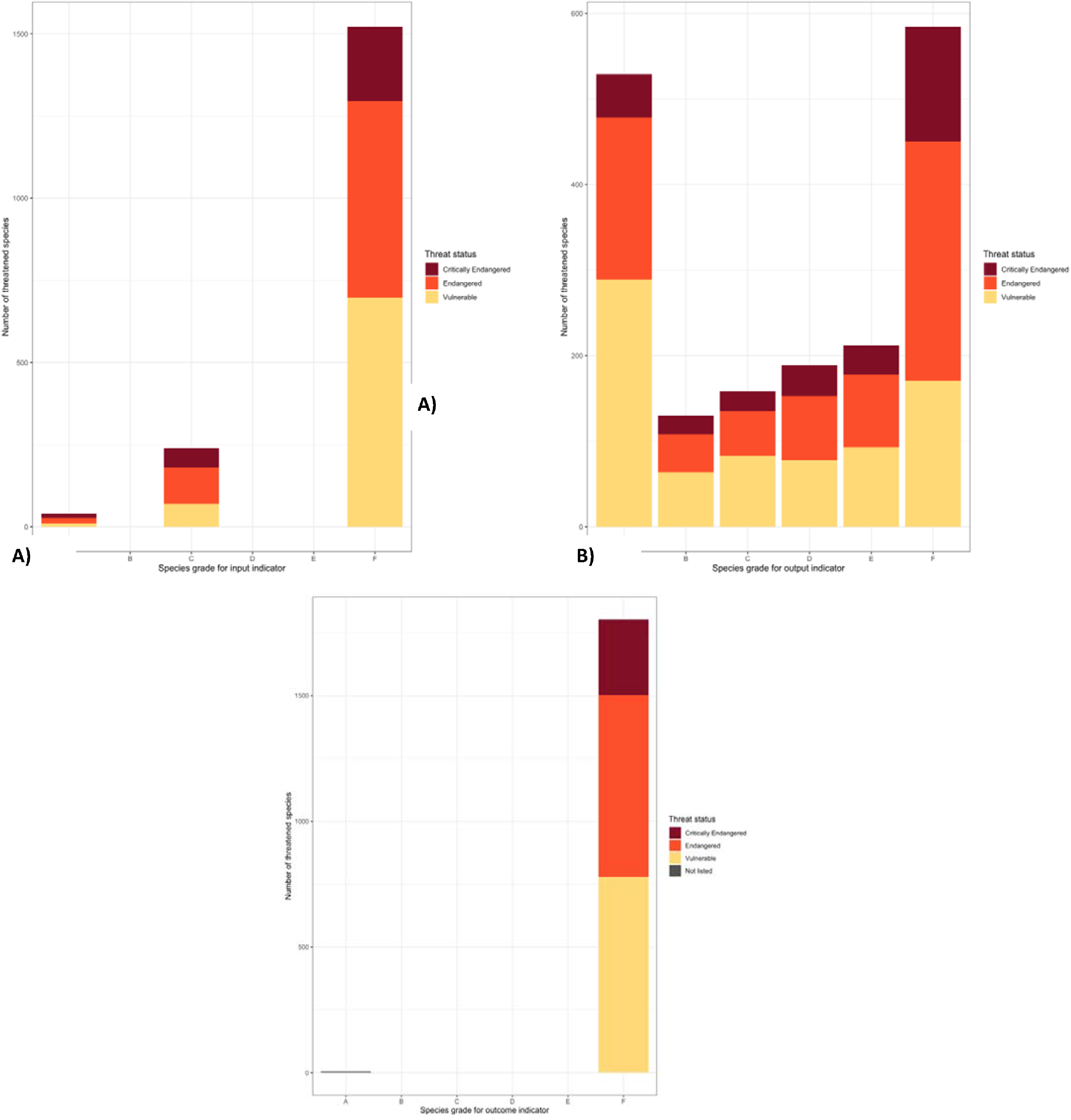
Input indicators (**A**), output indicator (**B**), and outcome indicator (**C**) across every threatened species (Y-axis) to provide an overall grade (X-axis). Critically Endangered species are highlighted maroon, Endangered species are highlighted orange, Vulnerable species are highlighted yellow, and Not listed species highlighted as grey.

Australia is failing in the recovery of threatened species. Australia scored an F grade on the input indicators (i.e., average of proportional recovery plans, and federal funding; **Fig 4**). For the output indicator, Australia scored a D (or a score of 0.48), indicating that on average 48% of species are meeting their species-specific protection target. Australia scored a F grade (or a score of 0.003) for the outcome indicator (i.e., threat status improvement). When we analyse individual indicators, Australia scored particularly low in recovery plans (F or 0.10) and dedicated funding (F or 0.08).

**Figure 4.**
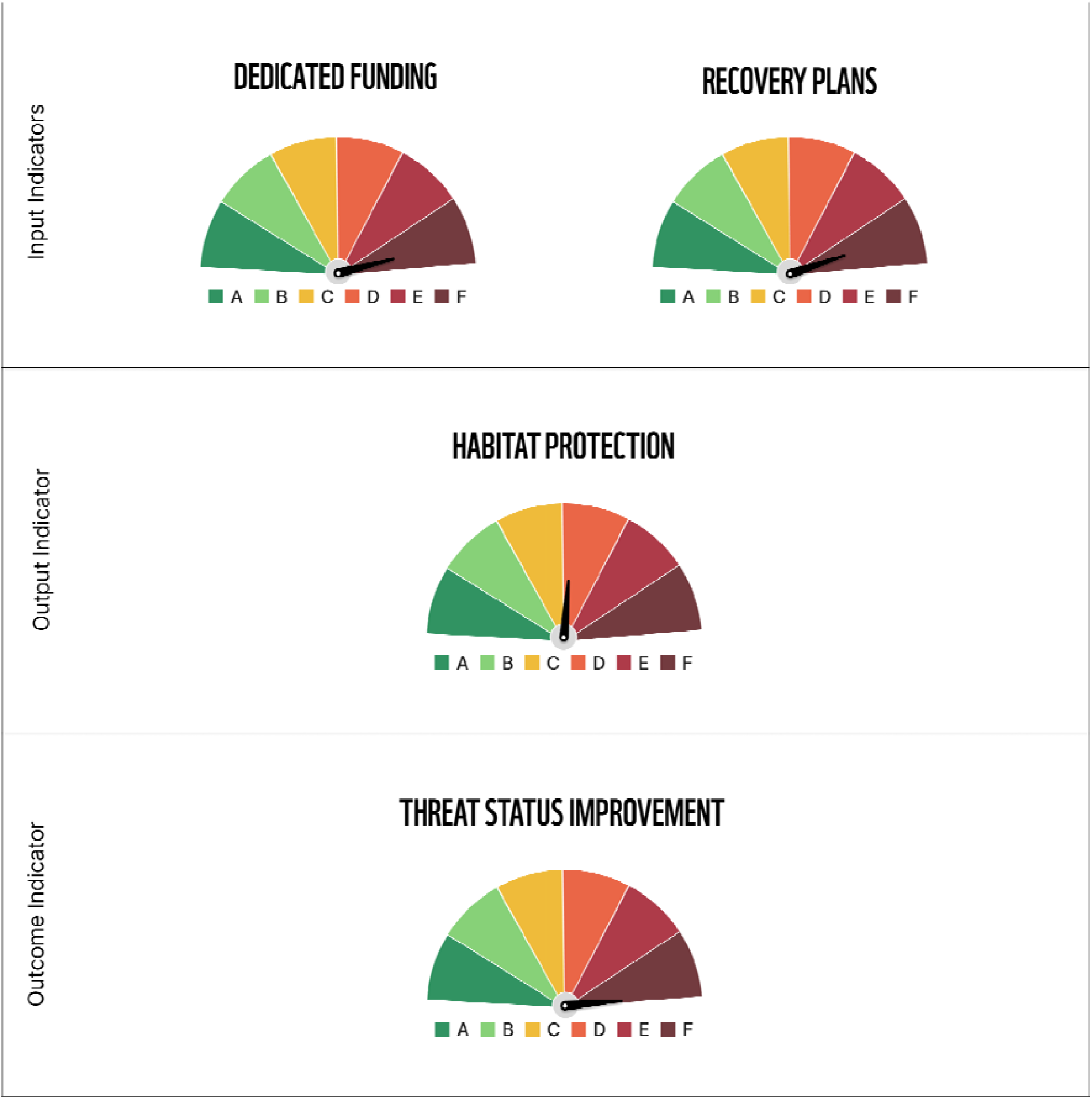
Threatened species recovery report card highlighting the results for the individual input indicators (top row), output indicator (centre row), and the outcome indicator (bottom row). The report card is broken up into six grades, F (dark red), E (red), D (orange), C (yellow), B (green), and A (dark green).

When we averaged the two input indicators evenly across electorates, our analysis shows that all electorates are failing, whereby one electorate achieved a D grade, 121 electorates achieved an E grade, and 29 achieved a F grade (**Fig. 5**). The electorates with the highest proportion of species with recovery plans were Braddon, Lyons, and Franklin, all of which are in Tasmania. The electorates with the highest proportion of species with funding were McMahon, Fowler, and Macarthur, all of which are in New South Wales. When we analysed electorates across the output indicator (i.e., protection), we found that 74 electorates achieved an A, 61 achieved a B, 15 achieved a C, and one achieved a D grade. The electorates with these high proportions of species with adequate protection were Lilley (Queensland), Petrie (Queensland), and Solomon (Northern Territory). When we analysed electorates across the outcome indicator, we found that all electorates are achieving a F grade, however the electorate with the highest proportion of species with improved threat status was Leichardt (Queensland). Durack (Western Australia) contains the species with the lowest proportions of species habitat protection, only one improvement in any species threat status, and the lowest proportion of species with any dedicated funding (averaging 6% of species).

**Figure 5.**
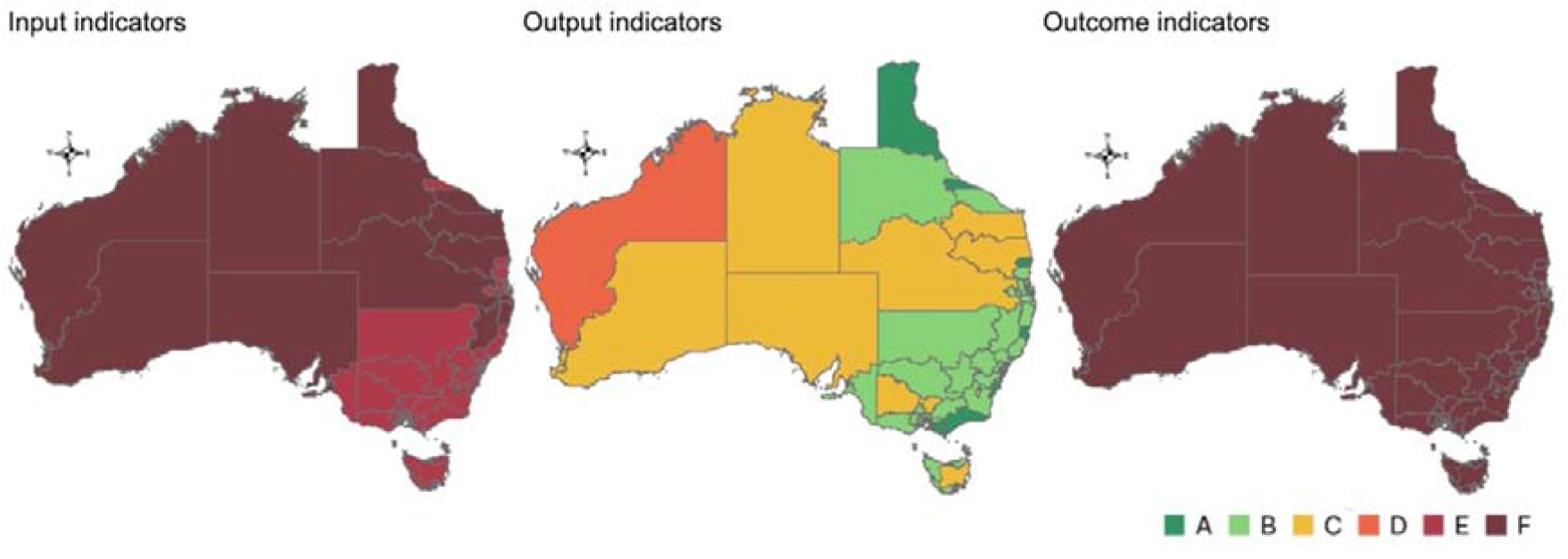
Maps of electorates highlighting both the input indicator grades (left), output indicator grades (centre), and the outcome indicator grades (right). There are six grades, F (dark red), E (red), D (orange), C (yellow), B (green), and A (dark green).

When we analysed the average grades of the input and output indicators across the different states and territories, we found that the Australian Capital Territory scored the highest in both (E grade and B grade, respectively). Queensland had the highest outcome indicator (F grade with a score of 0.008) as it had the most species that had improved in threat status (n=4 of the 5 species). When we analysed the average scores across all individual indicators, we found that the Australian Capital Territory contained species with the highest proportional dedicated funding (**Fig. 6**). Tasmania contained the species with the highest proportion of species with recovery plans. Western Australia had the least proportional species funding and habitat protection than all other states and territories.

**Figure 6.**
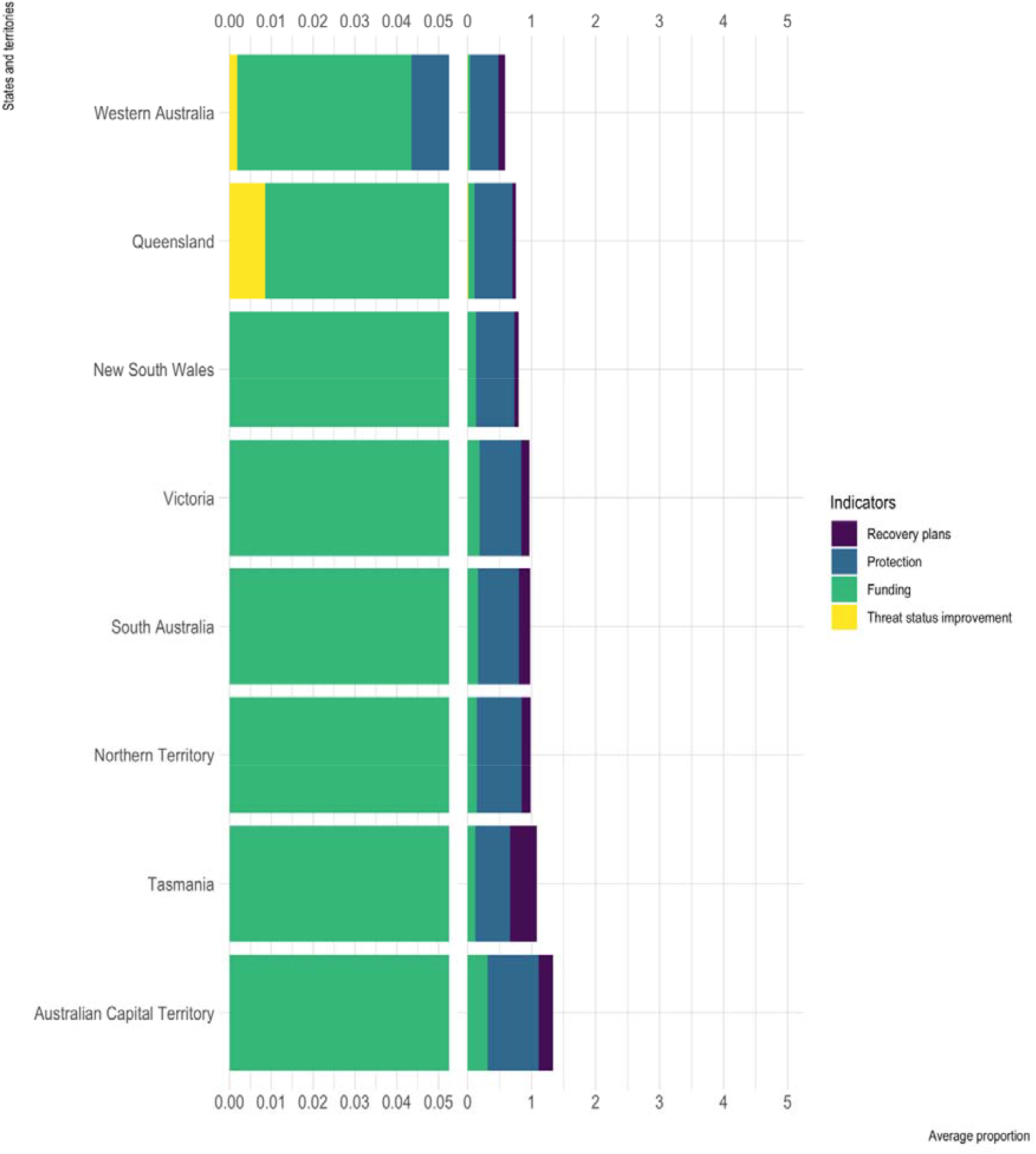
Bar chart showing the average proportion of indicators across Australia’s states and territories whereby funding is shown in green, habitat protection is shown in blue, recovery plans is shown in purple, and threat status improvement is shown in yellow.

Across the 547 local government areas (LGAs), 14 are achieving a D grade (including Ngaanyatjarraku (Western Australia), Karoonda East Murray (South Australia), and Albury (New South Wales)), 356 are achieving a E grade, and 177 are achieving a F grade for the averaged input indicator. When we analysed LGAs across the output indicator, we found 241 achieved an A grade, 230 achieved an B grade, 60 achieved a C grade, 14 achieved an D, and 2 achieved a E grade. The local government areas with the highest proportion of species with adequate habitat protection was Mount Magnet, Roxby Downs, Sandstone, Maralinga Tjarutja, Ngaanyatjarraku, Katherine, and Croydon (all received the highest score of 1). All LGAs achieved an F grade for the averaged outcome indicator, however the local government areas with the highest proportion of species with threat status improvement was Lockhart River (n=0.04). The local government areas with the highest proportion of species with recovery plans was Cocos Islands (n=0.67; **Fig. 7**). The local government areas with the highest proportion of species with funding was Ngaanyatjarraku (n=0.71 with 5 of 7 species with funding).

**Figure 7.**
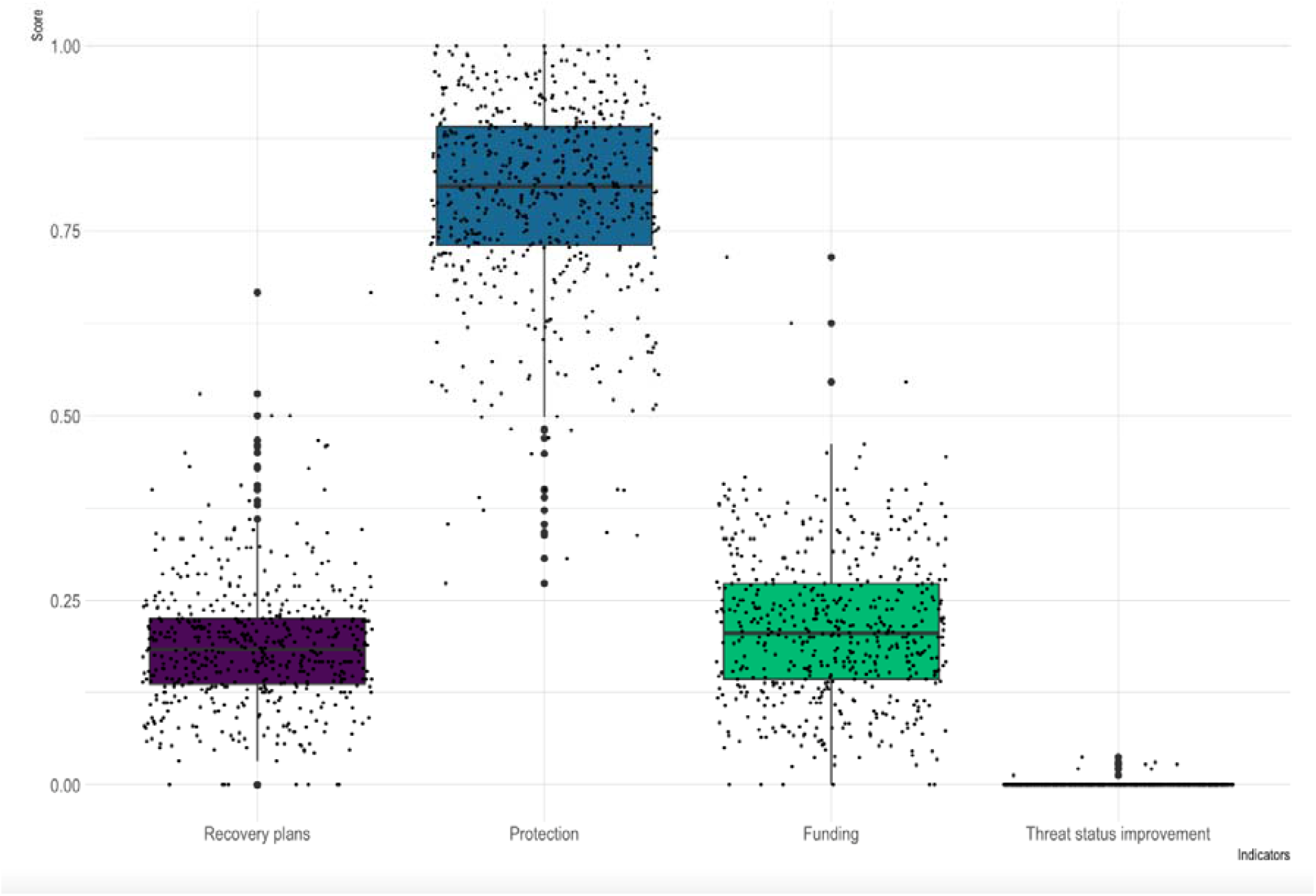
Box plots highlighting the variation between local government areas for recovery plans (shown in purple), habitat protection (shown in blue), funding (shown in green), and threat status improvement (shown in yellow).

## Discussion

Our threatened species recovery report card indicators were chosen to measure the current state of threatened species protection and recovery, in a systematic and repeatable way, given the information currently available. When implemented to Australia, it highlights the power of the result as it shows how poorly the nation is performing with just five species (0.3%) getting an A grade for outcomes, when compared to 1,521 species getting an F. When averaged across the nation, Australia gets an F grade for outcomes, as do all states and territories. Our results are potentially not surprising given the many recent reviews showcasing how nature laws are failing to protect Australian threatened species from extinction (Ward et al. 2019; Samuel 2020) but the nature of the report card highlights the dramatic shortfall in conservation attention threatened species are receiving.

While Australia is somewhat unique in that it holds an unusually high number of threatened species, we believe the broader methods used in this report card could be used in any country or site. The enormous increase in effort within the IUCN databases has meant that species and conservation information is increasingly available and national reporting efforts, often fostered through requirements like the SDGs and CBD, mean that nations are generating databases that can be utilised to populate the methods of a report card methodology (IPBES 2018; IUCN 2018; United Nations Environment Programme 2021).

We do note there are many indicators that could be added in the future to give a more complete picture of threatened species recovery. This comprehensive report card requires the Australian government to not only invest in more data collection but develop a national data infrastructure to support transparent reporting. One key missing indicator is habitat loss, which would be a key output measure. Habitat loss is not just the number one threat impacting the most threatened species in Australia (Ward et al. 2021) but across Earth (Maxwell et al. 2016). While there is an Australian legislative requirement to demonstrate the clearing of habitat will not have a ‘significant impact’ on the species recovery (Commonwealth of Australia 1999), we argue that all threatened species habitat is important to retain considering the downward trajectory towards extinction of these species (Ward et al. 2019). Unfortunately, there is currently no accurate data on vegetation loss at the continental scale.

As noted in the methods section, the accuracy of the funding recovery indicator is currently limited by a lack of data on federally funded threatened species conservation projects. Current publicly available data (Supplementary Table 1) does not specify the funding amount for all approved conservation projects. We could therefore only assess in a binary way whether a species was mentioned as receiving federal government funding or was not. Ideally this indicator should instead measure whether a species has received enough funding to ensure their recovery. Yong et al. (2022) developed cost models for threat abatement strategies across Australia, providing a framework that can account for spatial variables such as terrain and travel distance (Yong et al. 2022). Future report card analyses could use these models to estimate the approximate costs of threat abatement for each threatened species, and then assess the adequacy of provided funding using a similar metric to the habitat protection indicator. While this would provide a more realistic indicator of funding adequacy, it does not account for species-specific actions, e.g., translocation and ex-situ management, which are required to recover many threatened species (Bolam et al. 2022). Ideally, recommended improvements in monitoring, evaluation, and reporting under the EPBC Act (Samuel et al. 2021) should include a transparent system where recovery actions are identified for every threatened species, funded, implemented, and the outcomes monitored to ensure actions are effective and funding is sufficient.

Critical habitat conservation is another important consideration in recovering species. Critical habitat recognises threatened species’ requirements for high quality and well-managed habitat that, if lost or degraded, would likely result in the decline and/or extinction of the species (Camaclang et al. 2015). In Australia, under the EPBC Act, it is an offence to damage critical habitat that has been listed on the Register of Critical Habitat. However, critical habitat listing has only been listed for five species in the last 22 years. Critical habitat needs to be mapped, protected, and monitored by the Federal Government to prevent further human-induced extinctions.

In addition, a recovery team should be included within the set of indicators for species recovery. A recovery team is a collaboration of partners brought together by common objectives to develop and/or coordinate the implementation of a recovery plan, conservation advice or program for a threatened species or ecological community, or for multiple species or ecological communities. In Canada, recovery teams are appointed by the Minister and may focus on single species, multiple species, or whole taxonomic groups (Government of Canada 2021). It is only through a collaborative approach between government agencies, non-government organisations, First Nations peoples, scientists, industry and the broader community that the necessary management actions are likely to be effectively implemented to better protect and recover these species or communities. Recovery teams are one way to achieve collaboration and coordination in threatened species/ecological community management. In the future, we would like to include assessment of threatened species with a recovery team. Unfortunately, no comprehensive list of recovery teams exist, although the Australian government is working to create a national register of recovery teams and a framework for recovery teams to monitor and report on progress in achieving the objectives of recovery plans (Commonwealth of Australia 2022b).

Another key consideration for threatened species recovery is effective environmental legislation and policy. To ensure that federal, state and local legislation is effectively meeting environmental objectives, we require timely, independent reviews. Unfortunately, these reviews do not occur across all states and territories, nor at the local level, therefore, have been excluded from this analysis. Future iterations of the report card require quantitative analysis of not only legislation that prevent further decline but also those that allow positive environmental actions to prosper.

Overall, our threatened report card methodology shows Australia is failing in its efforts to recover threatened species. While progress in species habitat protection could be argued as fair, recovery plans, transparency in the costing and funding of recovery actions, and improving threat status are almost non-existent across the nation. The threatened species recovery report card has highlighted a clear failure of federal environmental legislation and international commitments to recover biodiversity. Like others, who have found threatened species policy and management ineffective (McDonald et al. 2015; Scheele et al. 2019; Ward et al. 2019), and that government spending is insufficient (Mccarthy et al. 2012; Wintle et al. 2019), we strongly encourage the Australian Government to take a critical leadership role in addressing the species extinction crisis by increasing habitat and vegetation protection, developing recovery plans for all threatened species, financing adequate recovery of such plans, and verification of recovery through monitoring and evaluation of species trajectories. Without these critical changes, we will leave a tragic legacy of extinction and fail our obligations to future generations of Australians, and the international community.

## Acknowledgements

We thank Kash Gunaretnam for their support in assisting with the design of some of the figures and Josie Carwardine for her revisions on earlier drafts.

